# Functional Exploration of Copy Number Alterations in a *Drosophila* Model of Triple Negative Breast Cancer

**DOI:** 10.1101/2020.10.18.343848

**Authors:** Jennifer E. L. Diaz, Vanessa Barcessat, Christian Bahamon, Ross L. Cagan

## Abstract

Accounting for 10-20% of breast cancer cases, TNBC is associated with a disproportionate number of breast cancer deaths. Despite recent progress, many patients fail to respond to current targeted therapies. Responses to chemotherapy are variable, and the tumor characteristics that determine response are poorly understood. One challenge in studying TNBC is its genomic profile: outside of *TP53* loss, most cases are characterized by copy number alterations (CNAs), making modeling the disease in whole animals challenging. We analyzed 186 previously identified CNA regions in breast cancer to rank genes within each region by likelihood of acting as a tumor driver. We characterized a *Drosophila* p53-Myc model of TNBC, demonstrating aspects of transformation. We then used this model to assess highly ranked genes, identifying 48 as functional drivers. To demonstrate the utility of this functional database, we combined six of these drivers with p53-Myc to generate six 3-hit genotypes. These 3-hit models showed increased aspects of transformation as well as resistance to the standard-of-care chemotherapeutic drug fluorouracil. Our work provides a functional database of CNA-associated TNBC drivers, and uses this database to support the model that increased genetic complexity leads to increased therapeutic resistance. Further, we provide a template for an integrated computational/whole animal approach to identify functional drivers of transformation and drug resistance within CNAs for other tumor types.

## Introduction

Breast cancer is the second most common cause of cancer deaths among women in the U.S.^1^. The most aggressive subtype, triple negative breast cancer (TNBC) makes up about 15% of breast cancers. TNBC is molecularly heterogeneous with few currently identified druggable molecular targets, poor therapeutic response, and low survival rates^2,3^. Less than 40% of women with metastatic TNBC survive 5 years^2^. Standard-of-care treatment of TNBC is limited to chemotherapy and, for some patients, immunotherapy including atezolizumab^4^. Meanwhile, advances in sequencing technology have opened new opportunities for understanding the mechanisms of tumorigenesis and drug response^5,6^. Such studies may improve breast cancer survival by improving predictions of progression in individual patients, identifying novel therapeutic targets, and improving the utility of our preclinical models.

Developing genetic models for TNBC is challenging. Most TNBCs contain mutations in *TP53*, however no other genes are commonly mutated^7^. Instead, computational work^7–9^ has identified extensive genomic copy number alterations (CNAs) often at overlapping sites between patients. For example, the *MYC* locus—a key regulator of basal-like tumor biology^7,10^—is commonly amplified in TNBC. Understanding the role of CNAs in TNBC would benefit from a comprehensive functional study of putative driver genes and their interactions in a whole animal context.

Recently, the *Drosophila* field has developed multigenic models of cancer to capture aspects of a tumor’s complexity; these models have been used to explore drug response including as a screening platform to treat a cancer patient with resistant disease^11–17^. While flies lack orthologs of some human tissues, epithelial tissues in *Drosophila* such as the eye and wing have proven useful for modeling cancer networks and identifying candidate therapeutics^11,15,17–22^.

In this study, we establish a *Drosophila* p53-Myc platform as a tool for exploring TNBC genomic complexity. To leverage this platform, we first used a computational approach to rank genes within common TNBC CNA regions. We then used our *Drosophila* p53-Myc platform to assess the functional relevance of many of the most highly ranked gene candidates within each CNA region based on their ability to enhance transformation in a whole animal platform. The result is a functional database of TNBC drivers within common CNAs. Finally, we used this database to build a library of more complex Drosophila TNBC models. In contrast to a p53-Myc model, these more complex lines failed to respond to fluorouracil—clinically relevant for TNBC—demonstrating that increased genetic complexity can lead to drug resistance and identifying candidate resistance factors. As an integrated approach, this work provides a path towards functionally deconvoluting the role of CNAs in tumor progression and drug response.

## Results

### *TP53* and *MYC* are the most common driver genes in TNBC

To determine the number of mutated driver loci in TNBC, we performed an analysis with the MutSigCV^23^ algorithm. We found 81% of TNBC tumors in the 2012 TCGA dataset^7^ include a mutation predicted to alter *TP53* function, consistent with the reported 80% of basal-like tumors^7^. No other driver genes were commonly mutated (Figure 1a; Supplementary Table 1). This is consistent with previous work showing that breast cancer is primarily driven by CNAs rather than point mutations^24^.

**Figure 1.**
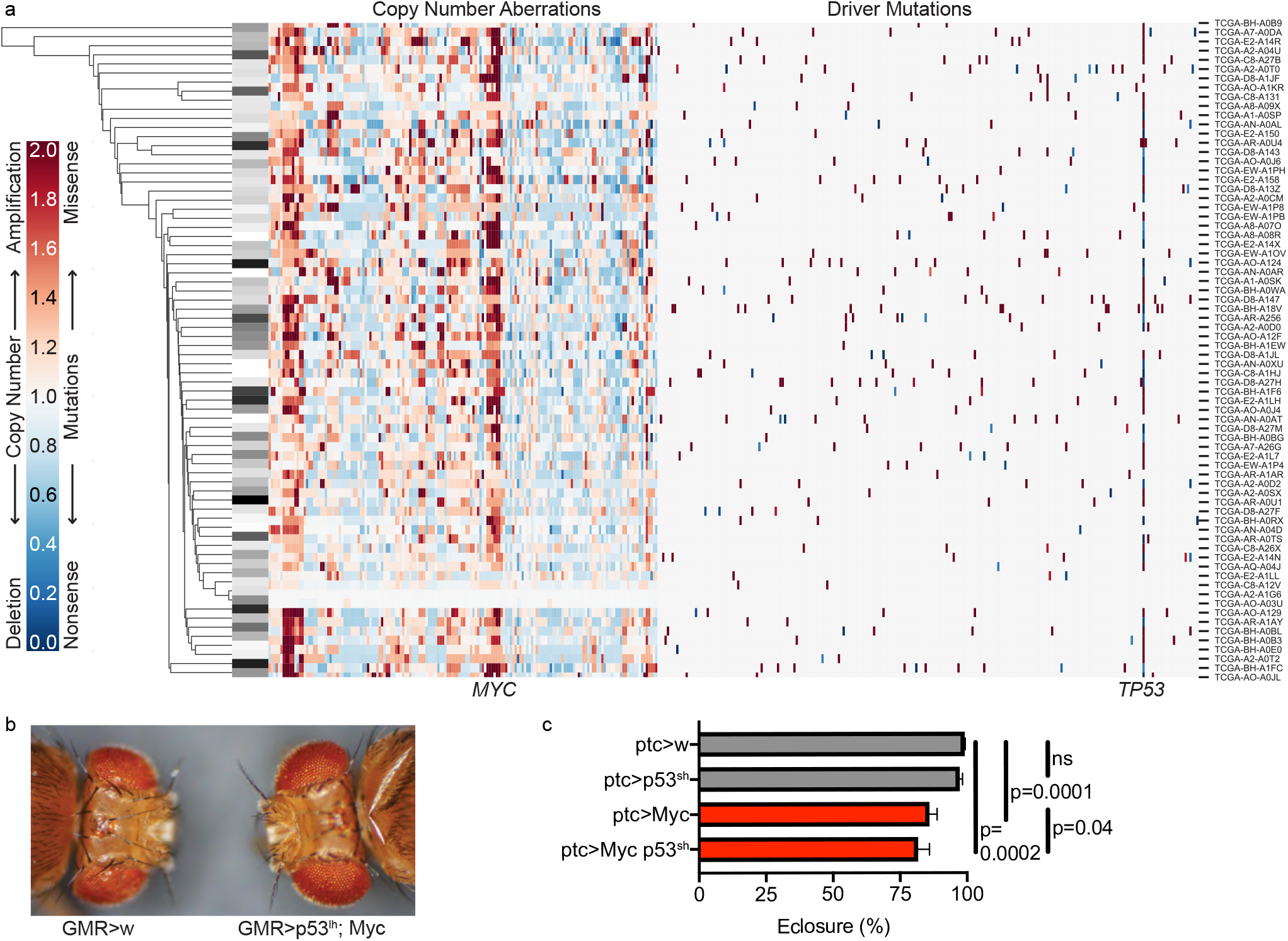
Altered *TP53* and *MYC* are present in TNBC tumors and reduced *Drosophila* survival. a) Hierarchical clustering of TNBC primary tumors in TCGA based on CNAs and mutations in putative driver genes. CNAs are shown in order of genomic location, and mutated putative driver genes are listed in alphabetical order. Overall survival of each patient is coded at left; white represents the longest and black the shortest survival. N=72. b) Fly eyes were enlarged when targeted by Myc overexpression plus p53 knockdown (*p53^lh^*; long hairpin). Genotypes in (b) are in the background of *yhsf* (see Methods). c) Quantification of survival of Myc-expressing flies in the presence *vs*. absence of p53 knockdown (*p53^sh^*; short hairpin). Kruskal-Wallis test: p<0.0001. N=14. Error bars represent SEM and do not reflect the paired nature of the data. P-values reflect Wilcoxon tests. See also Supplementary Figure 1 and Supplementary Tables 1.

The location and size of a CNA is not influenced solely by selective pressure, but also by chromatin architecture and the location of recombination hotspots^25–27^. As a result, a small portion of genes within a given CNA are responsible for the selective pressure and are therefore ‘drivers’, while neighboring genes are included due to proximity to the driver and mostly represent ‘passengers’ that do not appreciably contribute to disease progression.

As a first step towards exploring the genes within CNAs from the TCGA data, we began with 186 CNAs previously identified by GISTIC 2.0 and ISAR^7,9^. We performed hierarchical clustering on the 186 CNAs and driver gene mutations for the 72 primary TNBC tumors with both types of data available. We found that the tumors did not fall into discrete clusters, and tumors from patients with shorter or longer overall survival did not cluster together (Figure 1a). This suggests that in the aggregate, these genomic alterations do not explain differences in clinical outcome. This may be because some CNAs have more impact on outcomes than others, or because region-level analysis does not provide the resolution to understand the impact of individual driver genes. Therefore, a key step in using this genomic data to model TNBC is to identify drivers within the common CNAs.

The most common TNBC CNA was amplification of 8q24.1, a small region that—as identified by GISTIC 2.0—contains solely the oncogene *MYC* (Figure 1a). Excluding combinations of CNAs that overlap, the most frequently occurring combination of two events was mutation of *TP53* and amplification of *MYC*, in 72% of TNBC tumors. We therefore used the combination of these two genes, *TP53* and *MYC*, as the basis of a *Drosophila* platform designed to identify additional drivers.

### p53 and Myc promote cancer-like phenotypes in *Drosophila*

To characterize phenotypes produced by p53 and Myc, we generated individual fly lines containing trangenes that provide targeted expression of (i) *Drosophila* Myc (*UAS-Myc*) plus (ii) two different RNA-interference-mediated knockdown constructs targeting endogenous P53 (*UAS-p53^lh^* or *UAS-p53^sh^*; see Methods for details). Overexpression of Myc and strong loss of P53 (~80% in the presence of *UAS-Myc*) was confirmed by Western blot (Figures S1a, S1b). To confirm activity we used a *GMR-GAL4* driver to express *UAS-p53^lh^* and *UAS-Myc* in the developing eye field. *GMR>p53^lh^;Myc* flies exhibited enlarged eyes with normal ommatidial patterning (Figure 1b), presumably reflecting Myc-mediated cell enlargement.

We also tested the p53^sh^ and Myc transgenes individually and in combination using a *patched-Gal4* driver (*ptc*) that directs discreet expression of *UAS*-fused transgenes at several developmental stages. *ptc>Myc* survival to adulthood was reduced to 85.8%; *ptc>Myc,p53^sh^* exhibited further reduced survival to 81.5% (Figure 1c). Reducing P53 alone (*ptc>p53^sh^*) had no effect on survival. Similar results were obtained in the background of another fly strain (*yhsf*; see Methods), and with *p53^lh^* (Supplementary Figure 1c-d).

Using an inducible *UAS-GFP* to visualize transformed cells, we observed expansion of the *ptc* domain at the anterior/posterior boundary of developing *ptc>Myc* wing epithelia (‘wing discs’; Figures 2a, 2d, S2b). In contrast, the *ptc* domain was smaller with knockdown of p53 by *p53^sh^* (Figure 2a, 2d) and *p53^lh^* (Supplementary Figure 2b). In the presence of *ptc>Myc, p53^sh^* restored the *ptc* domain area to normal size (*ptc>Myc,p53^sh^*; Figure 2d). We have previously observed that p53 knockdown can reduce transformation phenotypes in the presence of an oncogene by reducing senescence^15^. Area reduction by *p53^lh^* was not significant (Supplementary Figure 2b).

**Figure 2.**
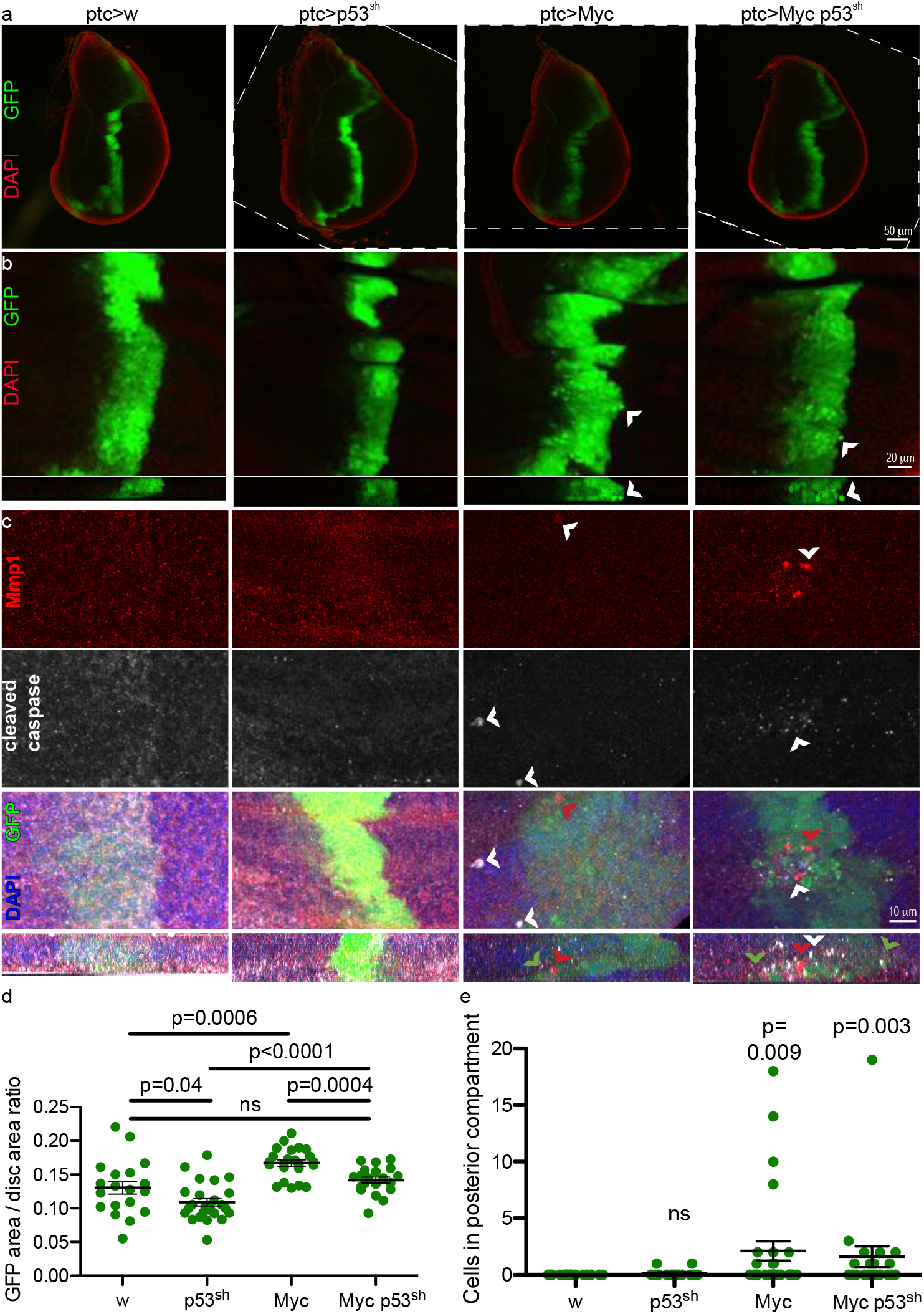
Overexpression of Myc promoted tissue expansion and cell migration in *Drosophila* wing discs. a) Representative wing discs of flies expressing combinations of *Myc* and *p53^h^*. Genotypically *white* (*w*) flies served as controls. DAPI in red highlights tissue boundary, GFP in green demarcates transgene expression. Some images rotated for comparison; borders indicated by dotted lines. b) Maximum projections of confocal stacks of the lower half of wing discs as in A. c) Maximum projections (upper three rows) and z-stacks (bottom row) of confocal stacks of the lower half of wing discs such as in (a) stained with Mmp1 antibody (red) and cleaved-caspase antibody (white), both indicative of cell migration29. Arrowheads mark delaminating cells; brightness and contrast were uniformly increased to improve visualization. In a-c: anterior at left, posterior at right, apical at top, basal at bottom. d) Quantification of transgenic tissue overgrowth produced by combinations of *Myc* and *p53^sh^* driven by *ptc-Gal4*. Kruskal-Wallis test: p<0.0001. p-values reflect student’s t tests. e) Quantification of cell migration in transgenic tissue produced by combinations of *Myc* and *p53^sh^* driven by *ptc-Gal4*. Kruskal-Wallis test: p=0.0075. p-values reflect Mann-Whitney tests compared to *w* controls. No significant difference was observed between *Myc* and *Myc,p53^sh^*. See also Supplementary Figure 2.

In confocal images of both *ptc>Myc* and *ptc>Myc,p53^sh^* wing epithelia, we observed transformed cells delaminating into the basal region of the epithelium (Figure 2b). These delaminating cells showed high levels of cleaved, activated caspase and matrix metalloproteinase (Figure 2c). Some of the delaminating cells were seen migrating away from the *ptc* domain (Figure 2e). Delaminating, caspase-positive cells (Supplementary Figure 2a) and migrating cells (Supplementary Figure 2c) were also seen with *ptc>p53^lh^;Myc*. The co-occurrence of caspase activation with migration is consistent with studies in other Drosophila cancer models^28–30^, in which migration was preceded by aspects of transformation and epithelial-to-mesenchymal transition. Based on the multiple aspects of transformation exhibited by our *ptc>Myc,p53^sh^* line, we concluded that it provides a useful genetic platform for identifying candidate functional drivers within regions of CNA.

### Prioritizing candidate driver genes from CNAs

Most of the 186 CNA regions identified by ISAR and GISTIC 2.0 contain dozens or hundreds of genes. Figure 3a provides a flowchart to summarize our approach for prioritizing these genes for functional testing. To curate likely driver genes, we first eliminated genes that (i) occur in known, common copy number variants, (ii) were not differentially expressed, or (iii) were associated with an increase in expression when copy number was reduced. To identify genes that were specifically relevant to TNBC, we analyzed the candidate genes for significance within the TNBC subset using a mild probabilistic cutoff. These steps reduced an original list of 12,621 candidates to 6,694 (see Methods; Figure 3a).

**Figure 3.**
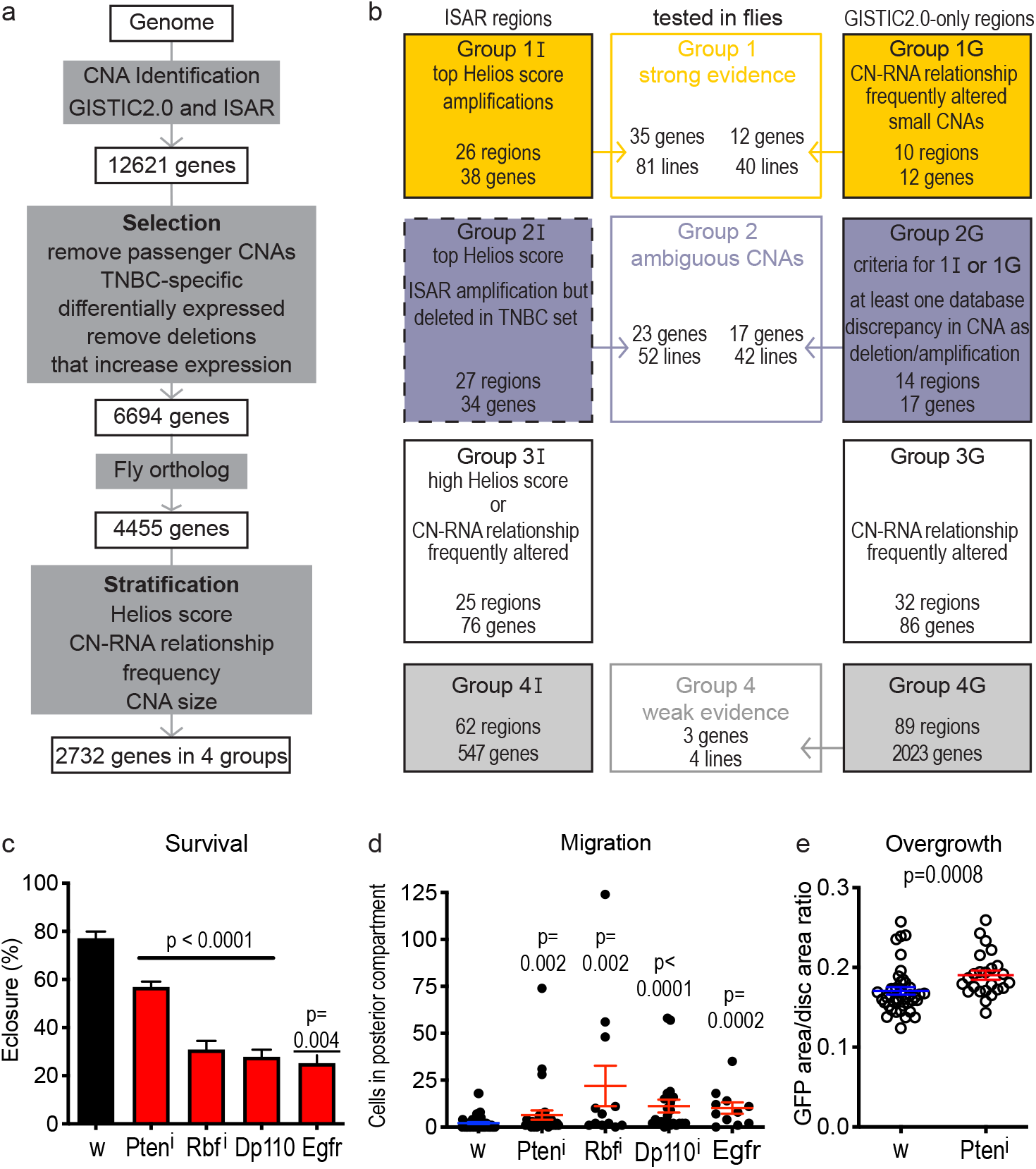
An integrated computational-functional screen to assess potential driver genes. a) Prioritization scheme of potential driver genes from CNAs based on TCGA data. b) Prioritized groups of genes for functional testing. Computational evidence is weaker for *Group 2I* (dashed border) than *Group 2G*. (c-e) Validating the screen: reducing activity by RNA-interference (Pten^i^, Rbf^i^) or increasing by over-expressing (Dp110, Egfr) four known driver genes in *trans* to *ptc>Myc,p53^sh^* led to decreased viability (c; n=4 for EGFR, n=8 otherwise), increased cell migration (d), and increased overgrowth of transgenic tissue (Pten knockdown shown as example). P-values reflect a student’s t test where data are normally distributed, or a Mann-Whitney test otherwise, compared to *w*. See also Supplementary Figure 3 and Supplementary Table 2.

When analyzing the TNBC dataset at this step, we found that some genes were deleted more frequently in TNBC than they are amplified, even though the CNA was identified as an amplification by ISAR or GISTIC 2.0, and vice versa (see Methods). We examined these genes separately in functional testing (below) as *Group 2*. Indeed, several cancer genes have recently been found to have paradoxical, context-dependent roles^31–33^. Because this study focuses on TNBC, we assigned each gene a CNA type according to our TNBC-specific analysis.

Of the 6,694 CNA genes, 4,455 genes had a clear fly ortholog, meaning they were available for functional testing. The genes were further stratified into groups based on: (i) each gene’s Helios score (for genes identified by ISAR) resulting from machine learning on TCGA and functional data in breast cancer^9^, (ii) whether the gene’s copy number influenced its expression, (iii) whether the gene was altered more frequently than other genes in the region, and (iv) the size of the region (see Methods). The results of our analyses are a set of prioritized genes for each common TNBC CNA (Supplementary Table 2).

For our functional studies, we focused first on the top ISAR genes (*Group 1I*; Figure 3a), small GISTIC 2.0 regions (*Group 1G*; Figure 4a), select genes with ambiguous CNA classification (*Group 2*), and three lower-ranked genes for which we had fly lines on hand (*Group 4G*). We then tested *Drosophila* lines representing these genes in functional assays, in many cases testing multiple lines for each gene (Supplementary Figure 3a). Control lines included three with transgenes unrelated to human genes, and one p53 null allele (expected to have no effect in the presence of p53 RNAi knockdown). In general, we determined how to analyze genes based on their status in our TNBC-specific CNA analysis: deletions were treated as tumor suppressors and amplifications were treated as oncogenes.

**Figure 4.**
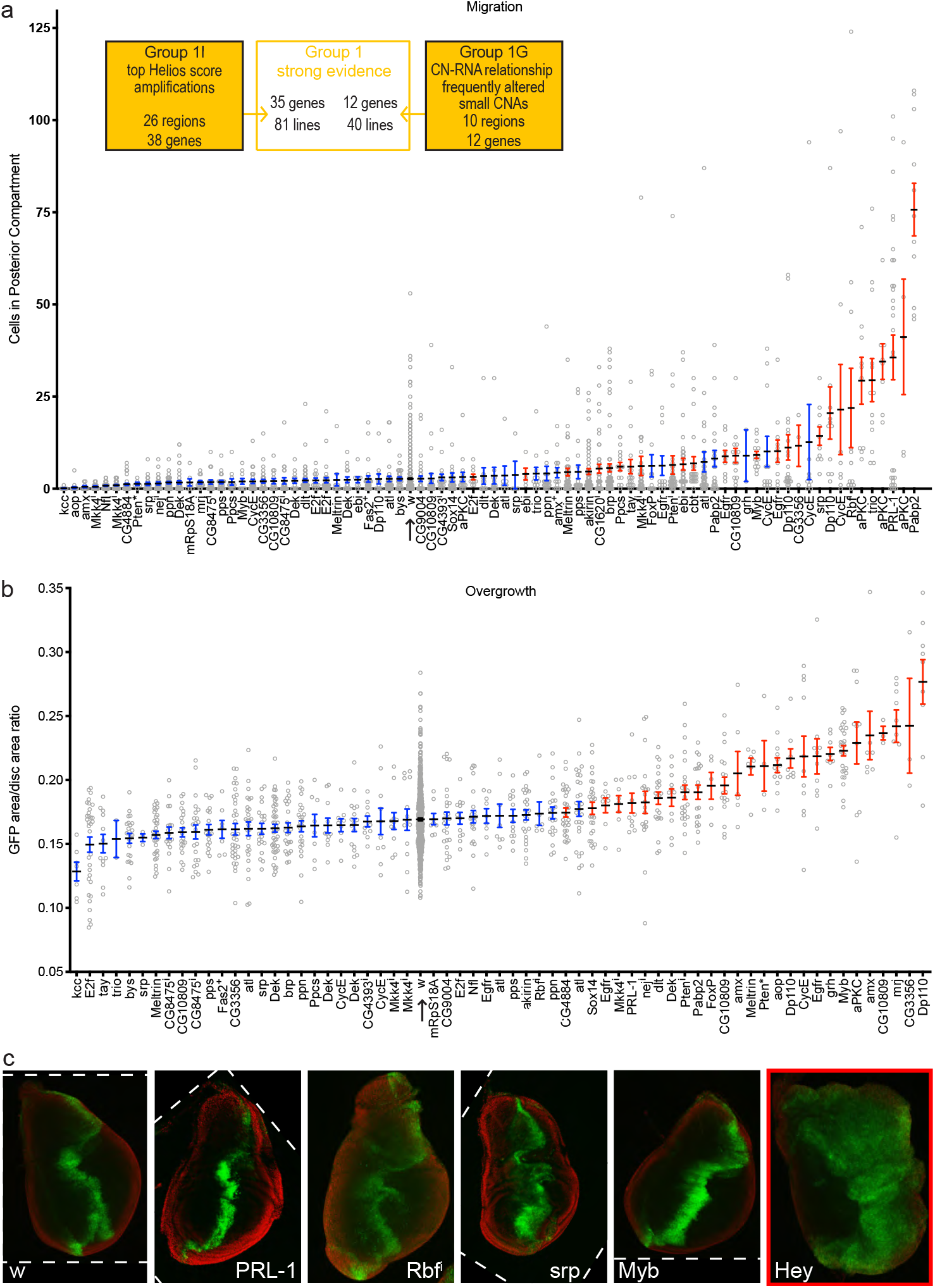
Driver genes produce tissue phenotypes in the background of *Myc* and *p53^sh^*. a) Quantification of cell migration for high priority genes. Altering genes marked in red directed a significant increase in migration compared to *w* (arrow), measured as p < 0.05 in the original experiment and false discovery rate (fdr) < 0.1 in this aggregate analysis. 16/52 from *Group1I* and 7/21 from *Group 1G* were significant. b) Quantification of transgenic tissue overgrowth for high priority genes. Altering genes marked in red directed a significant increase compared to *w* (arrow), measured as p < 0.05 in the original experiment and fdr < 0.1 in this aggregate analysis. Some genes that are significant in this figure were not significant in their respective experiments due to variation between experiments. i indicates RNA-interference mediated knockdown; * indicates a heterozygous null allele; + indications a duplication. 15/23 from *Group 1I* and 5/12 from *Group 1G* were significant. c) Selected phenotypes produced by specific driver genes: migration (PRL-1, Rbf^i^, srp), small overt mass (Rbf^i^), disruption of morphology (srp), transgenic tissue overgrowth (Myb, Hey), and large overt mass (Hey). Some images rotated for comparison; borders indicated by dotted lines. Migration and overgrowth were not quantified for Hey (marked with a red border) because the large overt mass phenotype was 100% penetrant. See also Supplementary Figure 4 and Supplementary Files 4-5.

### Driver genes enhance p53/Myc transformation phenotypes

Our computational analysis and ranking protocol identified a set of candidate drivers within TNBC-associated CNAs. To functionally assess candidates, we placed candidate transgenes *in trans* to *ptc>Myc,p53^sh^*, creating a ‘3-hit’ model by standard genetic crossing. Amplified genes were assessed by overexpression (*ptc>Myc,p53^sh^/<u>UAS-candidate</u>*); deleted genes were assessed by RNA-interference mediated knockdown (*ptc>Myc,p53^sh^*/<u>*UAS-RNAi*[*candidate*])</u> or by removing one functional copy (*ptc>Myc,p53^sh^/<u>mutant</u>^-/+^*).

To validate our approach, we tested four well-known tumor drivers: oncogenes Dp110 and EGFR were overexpressed, and tumor suppressors Pten and Rb were reduced by knockdown in the context of *ptc>Myc,p53^sh^*. In each case (4/4), introducing the driver gene decreased pupal survival (eclosure rate; see Methods; Figure 3c); adding Pten also decreased larval survival (pupariation rate; Supplementary Figure 3b). These genes also significantly increased migration of cells within the wing disc (Figure 3d). For Pten, an example of a driver that showed only mild migration, overgrowth of transgenic tissue was significantly increased (Figure 3e). In contrast, 0/4 *Group 4* lines and 1/4 control lines showed significant reduction in eclosure (Supplementary Figure 3c). Three lines of *Group 4* and three control lines showed significant reduction in pupariation, perhaps due to variation in genetic background (Supplementary Figure 3d). However, 0/4 *Group 4* lines and 0/4 control lines showed significant change in cell migration (Supplementary Figure 3e). Furthermore, 0/3 *Group 4* lines and 0/2 control lines showed significant increase in transgenic tissue overgrowth (Supplementary Figure 3f). In summary, well-known drivers enhanced aspects of transformation in this model while non-drivers did not.

As a first step in identifying functional drivers within the CNAs, we functionally tested 222 fly lines, covering 47 of the 50 genes in *Group 1*, 40 of the 53 genes in *Group 2*, 3 of 2023 genes in *Group 4G*, and 3 control genes. Of the 222 lines tested over all groups, 100 showed consistent lethality when placed in *trans* to *ptc>Myc,p53^sh^*, defining 69 genes (Supplementary Files 1-2 and Supplementary Table 3). To determine whether genes that decrease survival also increase aspects of transformation, we tested 66 of these genes (multiple lines in some cases; Supplementary Files 1-3 and Supplementary Table 3) for changes in cell migration. For genes that did not show a significant change in migration in an initial trial, we also measured tissue overgrowth. Together, 48/66 genes (63/116 lines) significantly increased the migration (Figures 4a and S4a; Supplementary File 4) or overgrowth (Figures 4b and S4b; Supplementary File 5) phenotypes, and were judged to be drivers.

We identified some differences in the percentage of drivers identified (“hit rate”) within each computationally-defined group. Most tested *Group 1* genes were drivers, including 56% of genes in *Group 1I* and 75% of genes in *Group 1G*. The higher hit rate for *Group 1G* suggests that genes in small GISTIC 2.0 regions are especially likely to be drivers. In *Group 2G*, where at least one database was in agreement with our TNBC-specific analysis for each gene, 76% genes were drivers, a similarly high hit rate to *Group 1G*. However, only 38% genes tested in *Group 2I*, in regions identified as amplifications by ISAR but deletions by our TNBC-specific analysis, were drivers. This suggests that algorithms such as GISTIC 2.0 and ISAR are powerful discriminators of functional amplifications and deletions. That is, genes appearing in regions identified only as amplifications are generally unlikely to function as tumor suppressors. We opted not to test the remainder of the genes in *Group 2I*.

Altogether, our analyses identified 48 identified drivers, alongside *MYC*. These 49 genes define a functional set of CNA-associated putative drivers (Figure 5a; Table) and account for the observed copy number aberration of 66 partially-overlapping, computationally-defined regions (Supplementary Table 4). Of note, several of the drivers have been reported in *in silico* or *in vitro* screens for drivers (Table; Supplementary Table 5)^24,34–38^. Furthermore, 4 of the genes were determined to have been mutated at a significant rate (Supplementary Table 1). Finally, we assessed these genes for effects on survival in the TCGA breast cancer dataset^39^. Twelve genes were significantly associated with increased progression or decreased survival (Figure 5b) in patients (log-rank p<0.1; Supplementary Table 6). Most (31/49) of these genes showed evidence of confirmation of driver status by at least one of these methods (Table). This analysis demonstrates the strength of our computation/genetics approach to identifying functional drivers.

**Figure 5.**
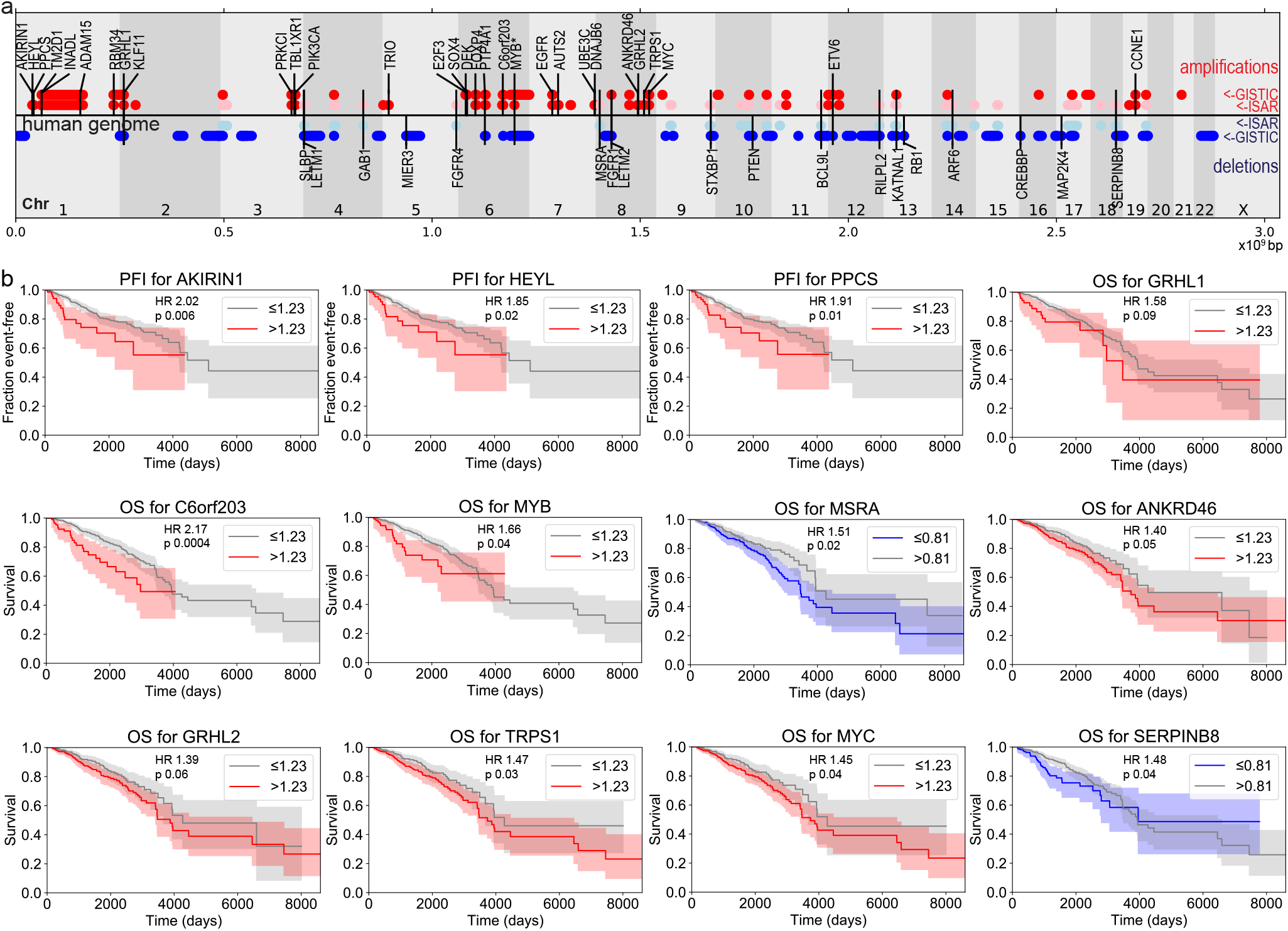
Well-known and novel driver genes in TNBC identified functionally. a) Map of genomic regions that are amplified (red) and deleted (blue) in TNBC, and location of functionally-validated driver genes that emerged in our screen. Group 2I regions, representing some ambiguity, are represented in pink and light blue. Genes with ambiguous CNA type are represented with a line extending through both amplified and deleted regions. Genes above the horizontal axis are oncogenes; genes below the axis are tumor suppressors. * indicates MYB can function as both. b) Kaplan-Meier curves of progression-free interval (PFI) or overall survival (OS) in the TCGA breast cancer dataset for CNA driver genes with log-rank p-value < 0.1. Amplified genes in red; deleted genes in blue. HR = Cox hazard ratio. See also Table and Supplementary Tables 4 and 6.

### Genetic complexity abrogates drug response in TNBC models

Work by a variety of laboratories including ours suggest that one source of resistance to therapeutic drugs is genetic complexity of the tumor. To test the effects of genetic complexity on drug response, we utilized our database of candidate CNA-associated drivers to generate six 3-hit models that reflect some of the genetic complexity in TNBC. Specifically, we paired *ptc>Myc,p53^sh^* with overexpression of oncogenes Dp110, Hey, Myb, Ppcs, or aPKC, or knockdown of the tumor suppressor Rop.

To test the effect of genetic complexity on drug response, we screened 72 FDA-approved cancer drugs, JQ1 (targeting Myc pathway activity), and five novel drugs that have shown activity in other Drosophila cancer models. Drugs were tested at 27°C by mixing into *ptc>Myc,p53^sh^* flies’ media and consumed orally. Fluorouracil, a drug used in the treatment of TNBC^40^, provided the strongest rescue of *ptc>Myc,p53^sh^*-induced lethality at both 27°C and 29°C (Figure 6a-c), mirroring activity in TNBC patients. In contrast, all six 3-hit TNBC lines built from our functional database failed to show significant rescue by fluorouracil at either 27°C (Figure 6d) and 29°C (Figure 6e). We conclude that, similar to other tumor types^15^, increased genetic complexity can lead to drug resistance in models of TNBC.

**Figure 6.**
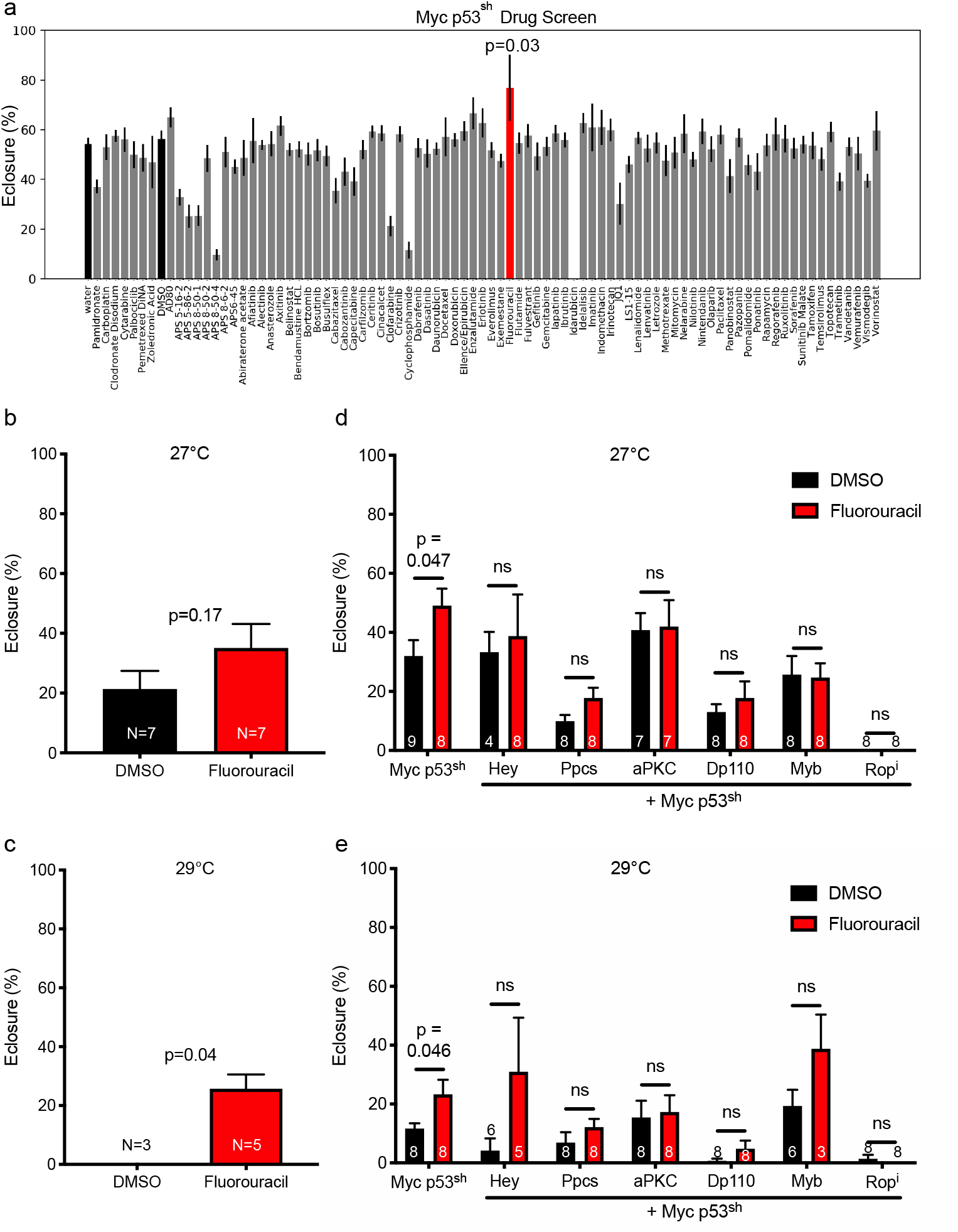
Genetic modifiers abrogate the response of p53-RNAi Myc to fluorouracil. a) *ptc>Myc* and *ptc>Myc,p53^sh^* strains were fed food containing screening-optimized doses of cancer drugs at 27°C; viability was assessed by survival to adult eclosure. Only fluorouracil significantly improved viability (Mann-Whitney U test *vs*. DMSO: p=0.03). b-c) Fluorouracil was tested on the *ptc>Myc,p53^sh^* line at (b) 27° C and (c) 29° C, and *ptc>Myc,p53^sh^* plus six selected driver genes at (d) 27° C and (e) 29° C. In each case tested, addition of an additional driver led to loss of fluorouracil-mediated rescue. Two-way ANOVA results were (d) genotype: p<0.0001, drug: ns, interaction: ns; (e) genotype: p<0.0001, drug: p=0.001, interaction: ns. Displayed p-values reflect t tests (see Methods). The final N for each condition is shown on each bar. See also Supplementary Table 7.

## Discussion

Producing useful genetic models of cancer and designing targeted therapies require an understanding of the genes driving tumor progression. This is especially challenging with TNBC, a disease in which most drivers emerge from copy number aberration rather than mutation, and carry many passengers with them. Approaches to identifying driver genes in this context have included statistical analyses of breast tumor sequencing data^34^, pan-cancer analyses^24,35,36^, cell culture screening approaches in transgenic lines^41^ and on breast cancer cell lines^37^, crowdsourcing^38^, and machine learning^9,42^. To our knowledge, this study represents the first systematic characterization of putative TNBC driver genes in a whole animal model.

Our data is consistent with previous evidence^7^ that TP53 mutation and MYC amplification are the two most common genetic aberrations in TNBC, and frequently occur in combination (Figure 1). In *Drosophila*, Myc promoted aspects of transformation including tissue overgrowth and cell migration (Figure 3). Knockdown of p53 enhanced Myc lethality, but not overgrowth or migration (Figure 2). This data indicates p53 likely affects other processes in the targeted tissue, such as senescence or metabolism, that can impact survival in Drosophila^15,43^. Our *Myc,p53^sh^* ‘base’ transgenic line constitutes a simple genetic animal model of TNBC. This model exhibited activation of matrix metalloprotease and caspase cleavage, previously validated markers of cell migration and metastasis-like behavior in fly cancer models^21,44,45^.

Many fly lines increased lethality when placed in *trans* to *ptc>Myc,p53^sh^*, including all known cancer drivers tested and several negative controls (Supplementary Figure 3c-d), suggesting that rescue of lethality provides a sensitive but not specific assay. However, only altering cancer-related genes—increasing ortholog expression of oncogenes or decreasing activity of tumor suppressors—altered transformation phenotypes in the *Drosophila* wing disc, providing a more specific second-line assay. Based on these functional assays, we identified or confirmed the cancer driver activity of 49 genes (Figure 5a). Some drivers such as *CCNE1* are well-established cancer drivers. Others such as *TRIO* are less well-studied but have previously appeared in the literature and databases of cancer drivers. *DNAJB6* has the highest Helios score in its region but was not considered a driver based on a cell culture assay^9^. Its observed effects in our overgrowth assay highlight the importance of testing for drivers in an intact epithelium where phenomena such as competition with wild-type tissue can be observed. To our knowledge, *PPCS, TM2D1, INADL, RBM34, C6orf203*, and *STXBP1* have not been linked to breast cancer. The variety of driver genes identified in this study underscores the potential of an integrated computational/experimental approach.

The high hit rates in *Groups 1 and 2G* (ranging 57-76%) indicate a significant enrichment in driver genes based on computational work alone. None of the *Group 4* genes tested were identified as drivers. This suggests computational features can enrich for driver genes over the initial identification of CNA regions by GISTIC 2.0 and ISAR. Some genes exhibited paradoxical properties. A small number of genes were amplified in the TCGA data but demonstrated decreased expression. For one such gene, *MYB*, we tested for both tumor suppressor and oncogene activity and found that knockdown led to increased cell migration (Figure 4a), but overexpression led to tissue overgrowth (Figure 4b). Similarly, our studies help deconvolve genes in ‘ambiguous’ *Group 2* CNAs. For genes identified as amplified in one TNBC database and deleted in another, 76% exhibited driver effects when tested according to whichever CNA occurred more frequently in TNBC.

In contrast, genes in regions only identified as amplifications by ISAR but more frequently deleted in the TNBC set (Group 2I) were less likely to be drivers (38% hit rate). Our assays demonstrated that the drivers we identified in this group function as tumor suppressors, despite appearing in amplified regions (Supplementary File 2). One example, *MSRA*, has been found to be a tumor suppressor in other functional studies^46,47^. Together, these data suggest the same gene altered by different CNAs may become an oncogene or tumor suppressor depending on subtype and biological context.

Developing a functional database allowed us to then ask a question significant to development of therapeutics: Does genetic complexity alter drug response, if the ‘base’ set of common drivers is the same? We found that while fluorouracil rescued our *Myc,p53^sh^* base model, introducing an additional transgene—including examples of both oncogenes and tumor suppressors, and genes from both Group 1 and 2—abrogated this response. This suggests that increasing genetic complexity reduces treatment response, confirming findings from work in other tumor models^15^. Our data indicate that genetic complexity may play a role in the poor outcomes for TNBC patients^3^, and suggest genes that are candidates to mediate chemoresistance.

Using a functional approach, we have identified multiple new driver genes that help explain specific amplifications and deletions found in TNBC patients. Some of these genes are novel or poorly understood cancer drivers that merit further research into the specific roles they play in TNBC. Further, we provide data that some of these genes can mitigate the response of a *Myc,p53^sh^* model to the chemotherapeutic fluorouracil. Further understanding the functional impact of these driver genes on drug response may help guide prognosis and drug selection based on genotype, and suggest new avenues for therapeutic development.

## Methods

### Analysis of TCGA data

Data from the invasive breast carcinoma dataset were downloaded from the TCGA Data Portal^48^. The dataset contains 1100 cases.

The somatic mutation annotation file (MAF) for this dataset contains 771 cases and 14375 genes. We uploaded this file to the MutSigCV public server^23^ to retrieve predicted mutated driver genes for breast cancer. Per the authors’ recommendation^23^, genes with q < 0.1 are considered significant. We also considered genes from COSMIC^36^, the TCGA Pan-Cancer Analysis^24^, the Vogelstein dataset^49^, and CIViC as of April 2016^38^ to be driver genes that might be mutated. Mutations in any of these genes are shown in Figure 1a. Known predisposing germline variants were found in 52 cases^7^. Mutations in any of these 10 genes were included in Figure 1a.

The level 3 copy number SNP array was downloaded for all 1020 samples for which it was available. Candidate genes included those reported by GISTIC 2.0 (total amp, basal amp, total del, and basal del datasets;^7^ and ISAR (from the total dataset and the basal dataset, defined by the authors as ER/progesterone receptor negative;^9^. For each of these, any gene symbol synonyms were converted to a consensus gene symbol from HUGO^50^. For each of these genes, coordinates were retrieved from NCBI build GRCh37 (also known as hg19). Raw copy number values were retrieved from the SNP array files labelled “no_cnv”, in which germline copy number variants known to occur in the population had been removed by TCGA. Whenever a value in the SNP array, in which the data are on a log_2_ scale, represented a region covering the entire gene, the copy number for that gene was retrieved and converted to a linear scale:

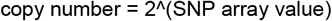

This value was then adjusted according to an estimate of tumor purity (the fraction of the sample that is tumor51. To account for germline variations in copy number, a fold change over the germline copy number was calculated, giving this formula for the final adjusted copy number:

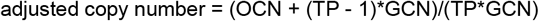

Where OCN = observed copy number in the primary tumor sample; GCN = germline copy number; and TP = tumor purity.

In the minority of cases where a tumor purity estimate or germline copy number was not available, a value of 1 was substituted for each of these parameters.

Thus, an adjusted copy number value of 1 represents normal in this study. Similarly to other studies of CNAs^34,52^, cutoffs of 2^0.3 and 2^-0.3 were used for amplifications and deletions, respectively. These represent gain and loss of one copy in approximately 40% of the tumor, respectively. The copy number for each region shown in Figure 1a represents the average of all genes in the region. The 72 TNBC cases with both copy number data and somatic mutation data are shown in Figure 1a.

Neither GISTIC 2.0 nor ISAR was applied to this dataset with TNBC specifically in mind. Thus genes that are not relevant to TNBC biology may appear in CNAs from these analyses, such as *ESR1*, which codes for ERα. GISTIC 2.0 was applied to the basal-like gene expression subtype, which only partially overlaps with TNBC. ISAR was applied to the ER/progesterone receptor negative subgroup that the authors termed ‘basal’, but this would also include CNAs relevant to HER2 positive tumors. These partially overlapping datasets resulted in some discrepancies. Some regions are called amplifications by GISTIC 2.0 in the total breast cancer set, but called deletions in the basal subtype set. Similar discrepancies also appeared between the ISAR and GISTIC 2.0 results (these were called “ambiguous CNAs” in this work). ISAR is explicitly designed to pick up amplifications even when they appear in the context of a larger deletion^9^, so this may explain some of the discrepancies.

To select genes in CNAs that are specifically relevant to TNBC biology and determine the direction of their effect, we applied an additional filter. Following Aure et al^34^, for each gene, we calculated the binomial probability of seeing the number of TNBC tumors with an amplification or deletion of that gene that appear, out of the 110 TNBC tumors in the dataset. We retained genes with a probability less than 0.05 for further analysis. Since this step represents an extra filter on top of an already rigorous CNA detection algorithm, we did not use multiple hypothesis correction. If more deletions of the gene appeared in the dataset than amplifications, we marked the gene as a deleted and treated it as a putative tumor suppressor; if more amplifications appeared than deletions, we treated it as amplified and a putative oncogene.

Next, we required that candidate genes in CNAs be differentially expressed. We downloaded the RNA-seq data for all samples for which it was available and extracted the rsem values for each gene. Following Akavia et al^52^, we retained genes with a standard deviation of greater than 0.25 for further analysis. We then converted the rsem values to z-scores for each gene. For each amplified gene, we performed a statistical test comparing the z-scores of primary tumors with that amplification and those without. We performed the equivalent analysis for deleted genes. Where the two distributions had equal variance, we used a student’s t test, and where they did not, we used a Welch test. All distributions were either normally distributed or had a sample size of at least 30, so we judged parametric statistics to be valid. We then applied a Bonferroni correction to all the resulting p-values.

We retained genes whose copy number had a direct positive relationship, or a non-significant relationship, with expression, for further analysis. Because amplifications could conceivably cause chromatin changes that reduce the expression of a tumor suppressor, we also retained genes that had reduced expression when amplified. However, as no theoretical mechanism could cause increased expression of a cancer driver gene when it is deleted, we removed genes that were associated with increased expression when deleted from further analysis.

For genes appearing in the ISAR dataset, we selected the top 1-3 genes by Helios score for *Group 1I* (Supplementary Table 2). The remaining genes in the top 3 (for regions with 12 or fewer genes) or top quartile (for regions with more than 12 genes) of Helios scores made up tier 2. Because the Helios score is based in part on the frequency of CNAs in the dataset as well as the relationship between copy number and expression, we did not directly consider these parameters to define *Group 1I*.

From the GISTIC 2.0 genes that did not appear in the ISAR data, we selected genes that 1) have a significant, positive relationship between copy number and expression, 2) appear in the top 3 (for regions with 12 or fewer genes) or top quartile (for regions with more than 12 genes) of genes in the region by the frequency of amplification or deletion occurring in the dataset, and 3) occurred in small (<10 genes) regions. These genes comprise *Group 1G* (Supplementary Table 2).

Genes marked for inclusion in *Group 1* were moved to *Group 2* if there were discrepancies between databases. In *Group 2G*, each gene belongs to at least one region identified by GISTIC 2.0, and some also belong to amplifications identified by ISAR. Genes in this group belong to at least one amplification and at least one deletion, and were analyzed as tumor suppressors or oncogenes according to the result of our TNBC-specific analysis. In *Group 1I*, genes belong to amplifications in ISAR but appeared to be deleted more frequently in our TNBC-specific analysis. Select genes in this group were analyzed as tumor suppressors. The remaining ISAR and GISTIC 2.0 genes comprise *Groups 3-4* (Figure 3b).

Finally, we converted all human genes in these 5 tiers to fly genes using homologs compiled from DROID^53^; downloaded 1/2014), DIOPT^54^; downloaded 10/2014), Homologene^55^, Ensembl^56^, and orthoDB^57^ (the latter three downloaded 7/2014). For genes with multiple possible fly orthologs, we performed the search manually in DIOPT and performed a tBLASTn^58,59^ search of the human protein sequence against the fly genome. The top scoring orthologs from each of these methods were used for testing (Supplementary File 3).

### Fly stocks

Experiments with overexpression of Myc used Bloomington stock 9675, with genotype hs-FLP y w; UAS-Myc. Initial experiments with p53 knockdown used a long hairpin under UAS control (Vienna Drosophila Resource Center stock 38235). This is represented as *p53^lh^*. However, although the long hairpin siRNA produced an effective knockdown, it also produced a faster migrating species on Western blot, possibly representing expression of a different P53 isoform. Due to this and the convenience of using a hairpin on the third chromosome, a short hairpin with guide sequence TGCTGAAGCAATAACCACCGA under UAS control on the third chromosome, generated in our lab, was used. We generated a recombinant third chromosome with this *UAS-p53^sh^* and *UAS-Myc*. This line has the genotype *w; UAS-Myc UAS-p53^sh^*. Both hairpins produce an approximately 80% knockdown in the presence of *UAS-Myc*, but *p53^sh^* does not exhibit the shift in band size (Supplementary Figure 2b).

In initial lethality experiments crossing *Myc p53^h^* to *ptc-Gal4*, low numbers of progeny resulted in survival variability after the first two technical replicates (flips). Therefore, only the first two technical replicates from each of seven different independent experiments (14 replicates total) were included in the analysis shown in Figure 1c.

We combined this line with patched-Gal4 on the second chromosome, the linked balancer *SM5(Gal80)-TM6B*, and a *hs-hid* construct on the Y chromosome to generate *(hs-FLP) (y) w /hs-hid; (ptc-Gal4; UAS-Myc UAS-p53^sh^)/SM5(Gal80)-TM6B*. This line was used for lethality experiments. We also generated a version with *UAS-GFP* on the second chromosome. *hs-FLP* was not detected in a PCR of fly genomic DNA from more than 30 flies from this line, indicating it likely lost the floating *hs-FLP* construct on the X chromosome, and so has genotype *(y) w /hs-hid; ptc-Gal4 UAS-GFP; UAS-Myc UAS-p53^sh^/SM5(Gal80)-TM6B*. This line was used for the overgrowth and cell migration assays.

To generate virgins of both these lines, bottles were incubated for 1-3 hours at 37°C twice between 1-5 days after lay, activating hid and resulting in death of all fertile males before eclosure. Virgins from heat shocks were only used for experiments and never put back into the stock.

Genotypes and stock numbers for lines used in the genetic screen are listed in Supplementary Files 1-2. Genotypes and stock numbers related to Figure 4a-b and Supplementary Figure 4 are in Supplementary Files 4–5. Stock numbers related to the genotypes shown in additional figures are as follows, in order (exclusive of *w*):

Figure 3c-e and Supplementary Figure 3b: VDRC35731, VDRC10696, in house UAS-Dp110 stock, BL9532

Supplementary Figure 3d (and related panels): BL20108, DGRC203601, BL17396, BL14265

Supplementary Figure 3e (and related panels): BL6659, BL6660, BL33623, BL6815

Figure 4c: BL20713, VDRC10696, d10248, F001574, F000566

Figure 6d-e: F00566, BL17464, F001672, in house UAS-Dp110 stock, F001574, BL28929

### Western blot

*765-Gal4* was crossed to lines carrying p53 knockdown or *UAS-Myc* to express the transgenes in the entire wing disc. Wing discs from L3 larvae were dissected in cold PBS. 10 discs were placed in RIPA buffer with protease inhibitor and phosphatase inhibitor. LDS sample buffer was added and the mixture was heated at 70°C for 5 minutes, then frozen at −80°C until further use. Samples were run on a Thermo Fisher 4-12% bis-tris NuPAGE gel per manufacturer instructions. Protein was transferred onto a PVDF membrane, which was then probed for p53 (DSHB p53-H3s 1:1000), syntaxin (DSHB 8B-3, 1:1000), or Myc (Santa Cruz Myc d1-717, 1:200) overnight at 4°C. Detection was performed with the Thermo Fisher Pierce™ ECL Western Blotting Substrate per manufacturer instructions.

### Lethality analyses

To measure survival with two or fewer genes of interest, *ptc-Gal4* was crossed to the genotype of interest (i.e. *UAS-Myc UAS-p53^sh^*) and the percentage eclosure rate was calculated as: 100 x [number of empty pupal cases/(number of uneclosed pupae + number of empty pupal cases)]. Crosses for the genetic screen were set up as follows at 25°C:

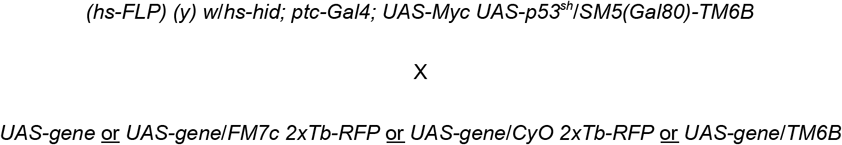

For genes of interest that were homozygous, or heterozygous on the first, second, or third chromosomes respectively^60^. For second and third chromosome genes of interest, males of the stock were used and virgins of the stock we generated as described above.

Because the chromosomes of the *SM5(Gal80)-TM6B* balancer are genetically linked by a reciprocal translocation, all progeny in the cross had either *ptc-Gal4; UAS-Myc UAS-p53^h^* or the balancer, which produces a tubby phenotype. Similarly, because of the balancers used, all progeny that did not have the gene of interest had a tubby phenotype. Eclosure was calculated as above only for the non-tubby pupae. Where the gene of interest was homozygous, a relative pupariation rate was also calculated as 100 x [number of *non-tubby* pupae/number of tubby pupae]. This was not calculated where the gene of interest was homozygous lethal and used with a balancer.

To account for the effect of variation in the genetic background of the stocks on survival, two biological replicates were set up for each gene of interest in each experiment. These were flipped four times, for a total of eight replicates for each genotype. Eclosure and relative pupariation for each gene of interest were compared to control with a student’s t test where distributions were normal or a Mann-Whitney u test where distributions were not normal. When either eclosure or relative pupariation was lower than control (p<0.05), the experiment was repeated. Genes that significantly lowered lethality in at least two independent experiments were considered candidates for tissue phenotype analysis. For simplicity, the aggregate of repeat experiments are shown in Figure 3c and S3b-d.

### Wing disc analyses

To visualize transformed tissue, we performed the equivalent crosses at 25°C as described above with a UAS-GFP construct on the second chromosome. Wing discs from non-tubby L3 larvae were dissected, fixed, and mounted in Vectashield with DAPI. Apical-basal and anterior-posterior axes were determined by locating the patched-expressing region of the peripodial membrane^61^. Cleaved caspase and matrix metalloprotease were visualized by immunostaining the fixed tissue (Cell Signaling antibody #9661, 1:200; DSHB antibody 3B8D12, 1:10). To assess migration, the discs were visualized at 40X magnification on a Leica DM5500Q confocal microscope and the number of GFP-expressing cells in the posterior compartment was counted.

Genes of interest that did not have a significant effect on migration were tested with the more laborious overgrowth assay. Each disc was imaged at 10X magnification, using identical settings within each imaging session, and the red and green channels were exported to separate tiff files. The outline of the disc was selected using the magnetic lasso tool in Photoshop, and the number of pixels in the enclosed areas was measured. The number of green pixels within the disc was measured in ImageJ using the threshold tool at the default setting and then the analyze particles tool. The relative amount of transformed tissue in each disc was then calculated as: number of green pixels in disc/number of pixels in disc.

For each of these two calculations (cell counts for migration, and disc area ratios for tissue overgrowth), the values for each set of discs of a genotype of interest were compared to control, using a student’s t test where distributions were normal or Mann-Whitney u test where distributions were not normal. For Figure 2 and Supplementary Figure 2, two-tailed statistics were used. For the genetic screen, only samples with a mean greater than control were tested, and one-tailed statistics were used. We then calculated the false discovery rate (FDR)-adjusted p-values using the Benjamini-Hochberg method available in the Python package Statsmodels^62^.

In the genetic screen, to account for variation in the baseline characteristics of the *ptc>Myc,p53^sh^* line over time (due to genetic drift) and slight variation in imaging settings from experiment to experiment, these statistical tests were first performed using only the controls from each individual experiment. However, as some experiments contained only a small number of samples for each genotype, and in order to calculate an FDR (above), we also performed the comparisons after aggregating the data for all experiments. Fly lines resulting in enhancement over control with p<0.05 in the individual experiment *and* FDR<0.1 in the aggregate were considered significant.

### Drugs

We used a library of drugs that are FDA-approved to treat cancer, and added several drugs that are typically used for TNBC, as well as novel compounds in the lab that had shown promise in other models. Doses were approximations of the maximum tolerable dose, based on previous experience with other fly models, with lower doses of some drugs based on preliminary experiments on p53/Myc (Supplementary Table 7). Drugs were dissolved in either water or DMSO, and then diluted 1:1000 by volume in the fly food. Water soluble drugs were compared to a ‘water’ control where no drug was added to the food. Drugs dissolved in DMSO were compared to food with 0.1% DMSO by volume.

### Drug Experiments

We crossed *ptc-Gal4* to *w; Myc,p53^sh^* at 27°C, where the rate of eclosure on DMSO was 56%, lower than at 25°C. Flies were allowed to lay on four replicates of fresh drug food for each drug for 24 hours. They were then flipped into another set of four replicates each and allowed to lay for 24 hours, giving a total of eight replicates for each drug condition. The percentage eclosure rate was calculated as: 100 x <u>number of empty pupal cases</u>/[number of uneclosed pupae + number of empty pupal cases]

The crosses for Figure 6b-e were performed similarly to those for the genetic screen. Virgin females of *hs-FLP/hs-hid; ptc-Gal4; Myc,p53^sh^/SM5(Gal80)-TM6B* were crossed to a control line with genotype *w* or genes of interest from the screen. The percentage eclosure rate was calculated as above only for the non-tubby pupae. The relative pupariation rate was also calculated as a percentage: 100 x [number of non-tubby pupae/number of tubby pupae]. Drug and food conditions were the same as above.

### Statistics

Hierarchical clustering was performed in Python 3 with the graphing package Seaborn^63^.

Statistical tests were performed either in Python 3 with the statistical packages Scipy^64^ and Statsmodels^62^ or Prism. In Figure 6d-e, the 2-way ANOVA and multiple t-tests (unpaired) functions in Prism 8 were used.

Survival analyses utilized the breast cancer portion of the TCGA Pan-Cancer clinical endpoints database^39^ and were performed using the built-in log-rank test, Cox proportional hazard model, and Kaplan-Meier curve function in the Python package Lifelines^65^.

## Acknowledgments

We thank the Cagan Laboratory for important discussions, and particularly Sindhura Gopinath and Chana Hecht for technical assistance. We thank Avi Ma’ayan, Giovanni Ciriello and Yasin Senbabaoglu for advice, Benjamin Neel for sharing data, and Dana Pe’er for sharing ISAR data. We thank Peter Smibert for designing key knockdown sequences, and Masahiro Sonoshita for generating and providing the P53 knockdown line. Funding: This work was supported by grants from the National Institutes of Health (U54 OD020353, R01 CA109730, T32 GM007280), and Mary Kay Ash. Competing interests: The authors declare that they have no competing interests. Data and materials availability: All data and accession numbers needed to evaluate the conclusions in the paper are present in the paper and/or the Supplementary Materials. Code is available at github.com/jennifereldiaz/fly-tnbc. Additional data related to this paper may be requested from the authors. Author contributions: Conceptualization, experimental design: JELD, RLC. Software, genetics, visualization, writing (initial draft): JELD. Drug screen: VB, JELD. Drug experiments: CB, JELD. Writing (reviewing and editing): RLC. All authors reviewed and approved the final version.

---

**Table.**
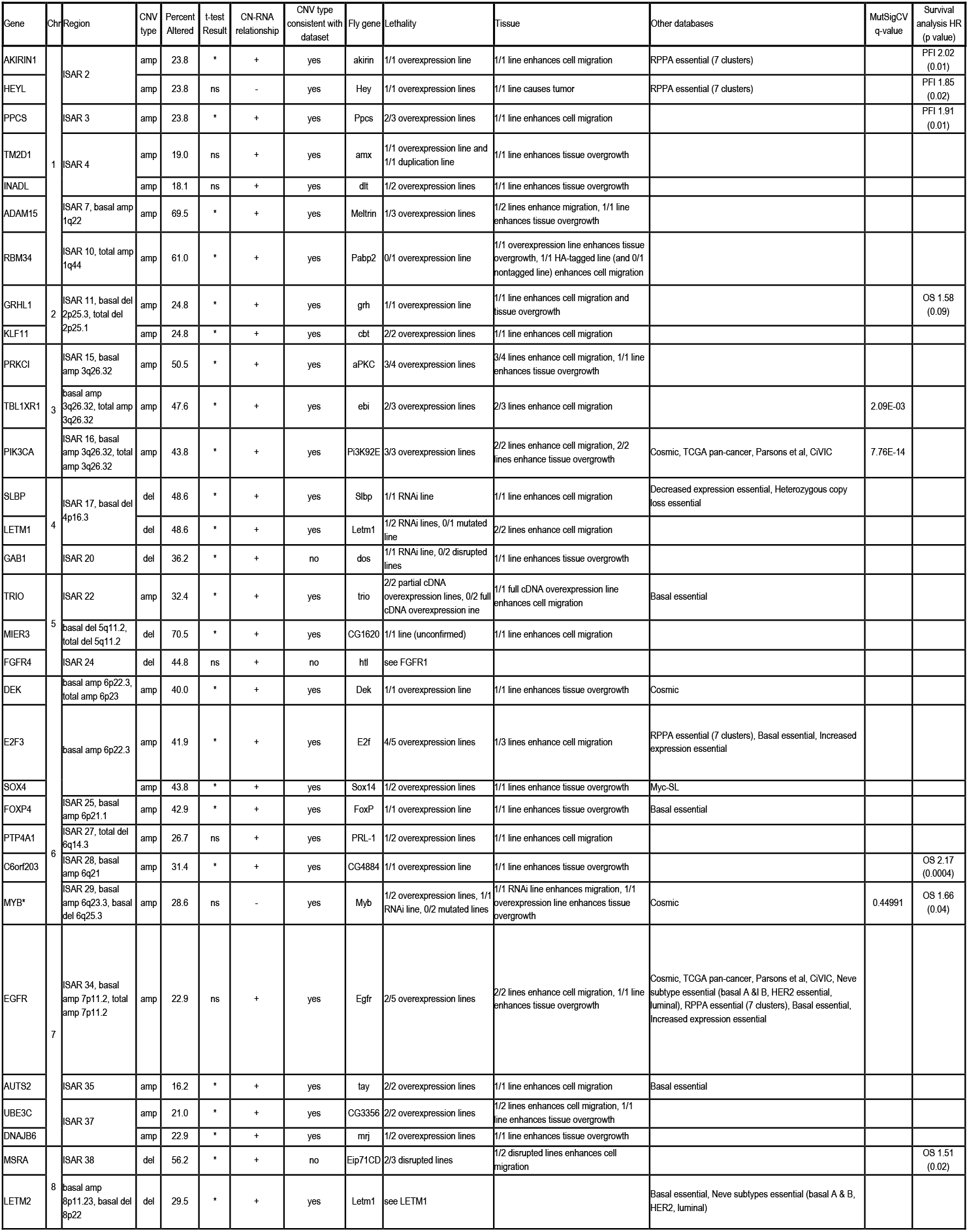
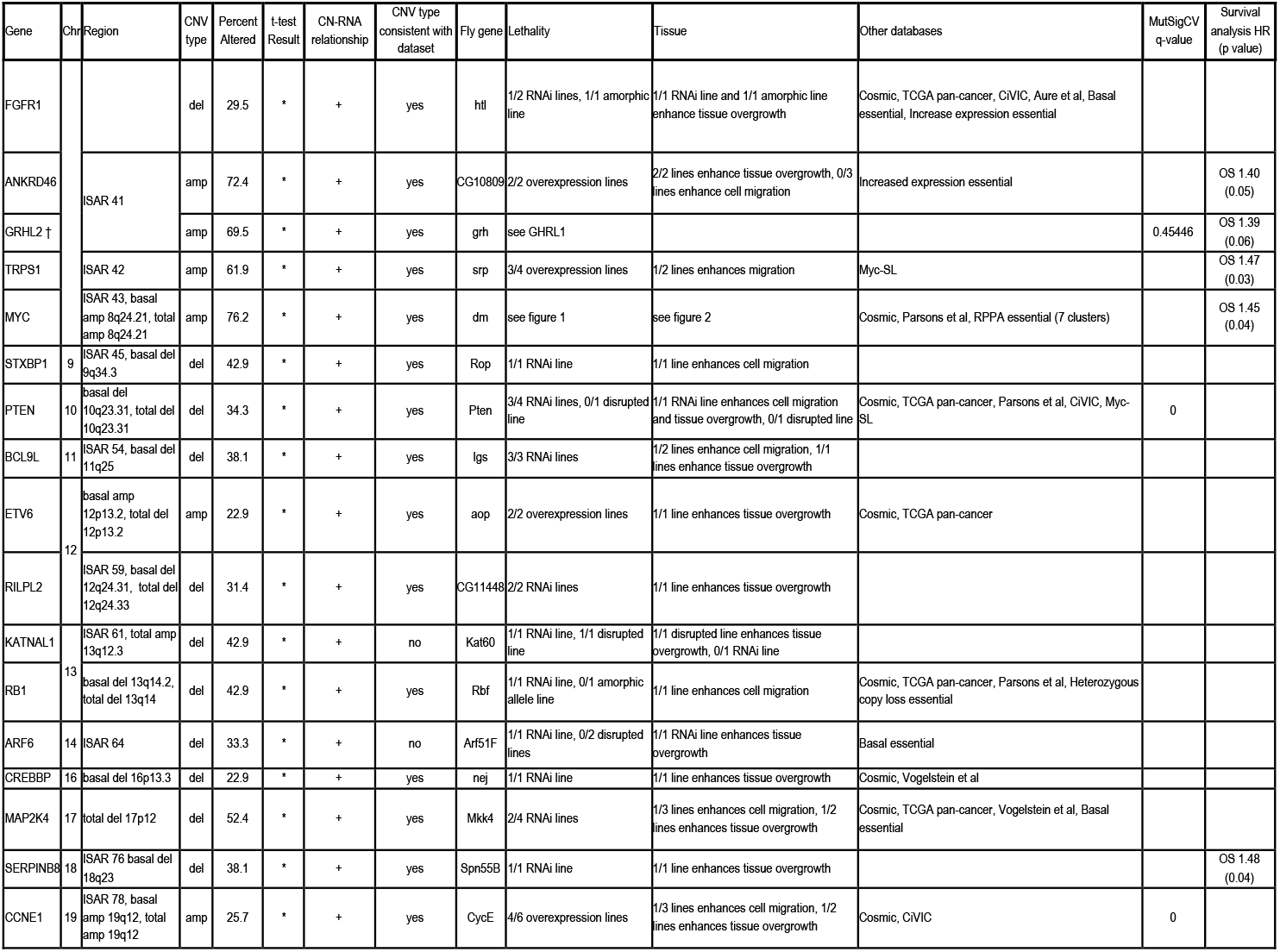
Computational and functional information on each driver gene. * indicates that MYB copy number has a possible negative relationship with expression (Supplementary Table 2) and was tested as both an oncogene and tumor suppressor. † indicates that GRHL2 did not meet criteria for inclusion in the screen but was tested because it shares an ortholog with GRHL1. MutSigCV q-value is shown only for genes with q < 0.1. Progression free interval (PFI) and overall survival (OS) p-values reflect a log-rank test for patients with the aberration vs. without. Only genes with p < 0.1 are shown, all of which the Kaplan-Meier curve plot indicates poorer prognosis with the CNA (Figure 5B-C). See also Supplementary Tables 1, 5, and 6.

## Supplemental Figure, Table, File Legends

**Supplementary Figure 1.**
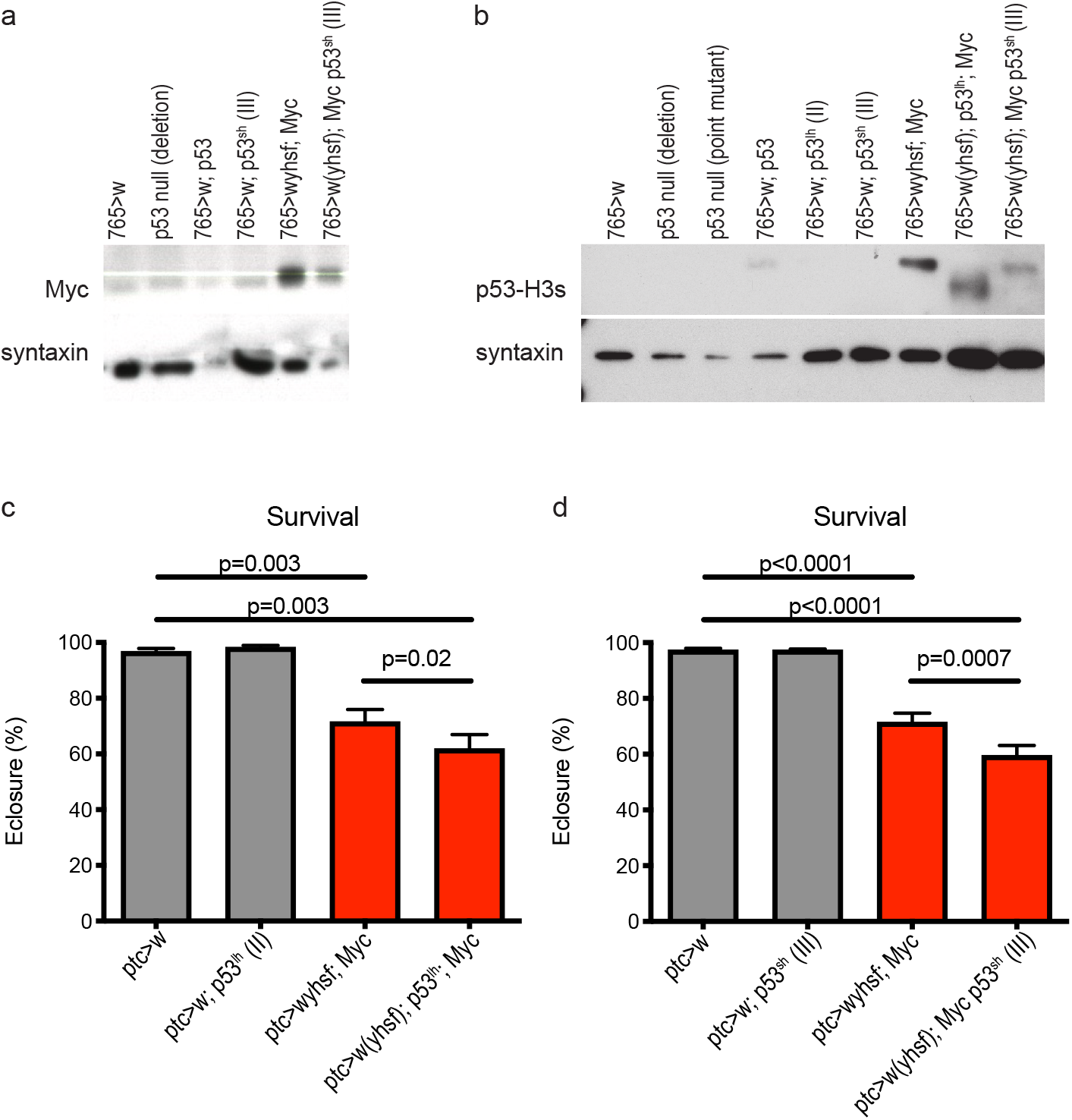
Myc and p53-RNAi synergize to impact survival in *Drosophila*. a) Western blot of Myc in fly wing discs. b) Western blot of p53 in fly wing discs. a-b were repeated in a separate experiment. c) Survival of *Myc*-expressing flies when combined with *p53^lh^*. d) Survival of *Myc*-expressing flies when combined with *p53^h^*. In c-d, Kruskal-Wallis test p<0.0001 and N=11. P-values reflect Wilcoxon tests. *y* and *hs-flip* elements were present on the X chromosome where denoted. Related to Figure 1.

**Supplementary Figure 2.**
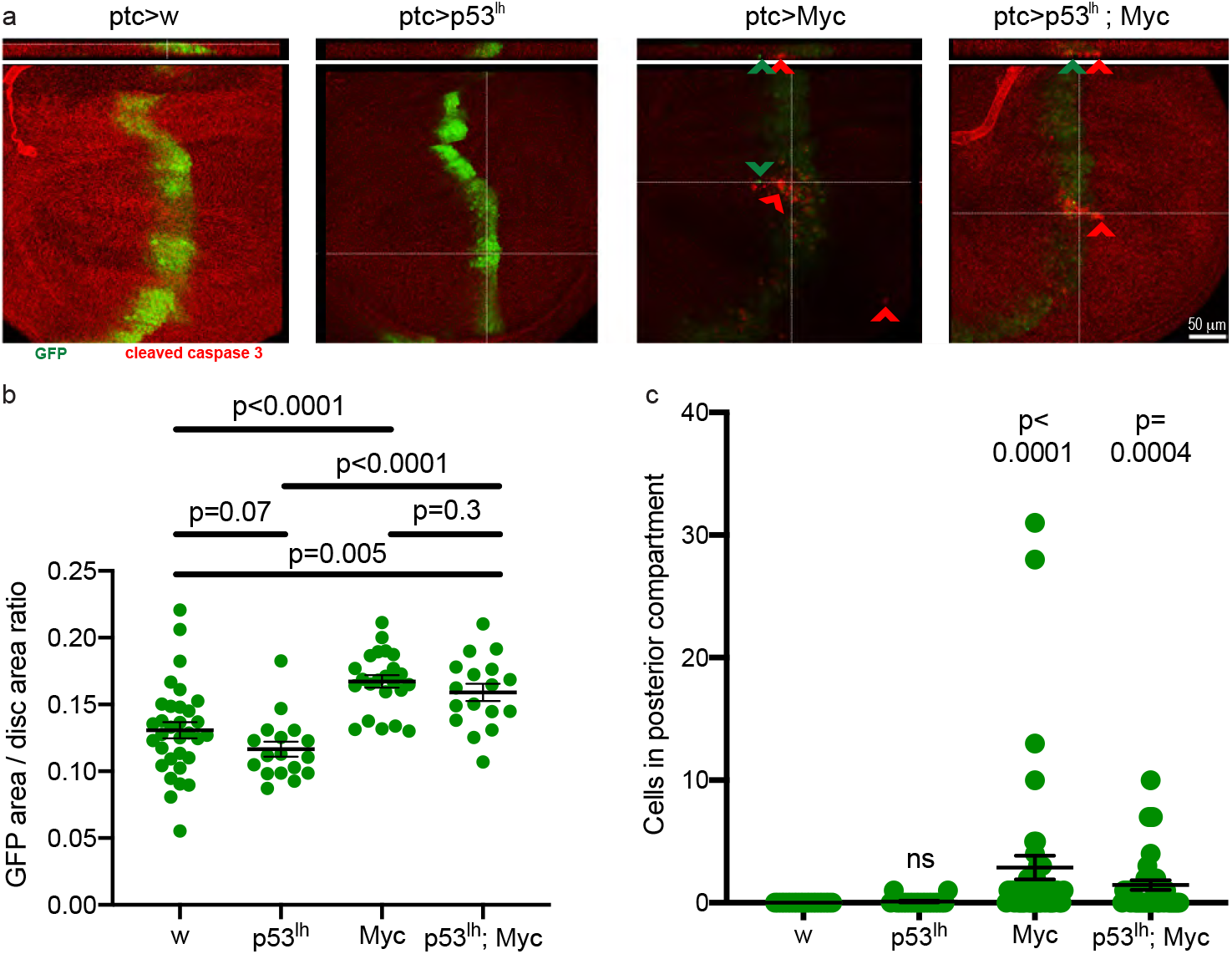
Overexpression of Myc induced tissue expansion and cell migration in *Drosophila* wing discs. a) Maximum projections (bottom) and z-stacks (top) of confocal stacks of the lower half of wing discs stained with a cleaved-caspase antibody (red). Arrowheads mark delaminating or migrating cells; brightness and contrast were uniformly increased to improve visualization of staining. Note *p53^lh^* was used rather than *p53^sh^* (figure 2). Magnification: 40X. Anterior at left, posterior at right, apical at top, basal at bottom. b) Quantification of transgenic tissue overgrowth produced by combinations of *Myc* and *p53^lh^* driven by *ptc-Gal4*. Kruskal-Wallis test: p<0.0001. P-values reflect student’s t tests. c) Quantification of cell migration in transgenic tissue produced by combinations of *Myc* and *p53^lh^* driven by *ptc-Gal4*. Kruskal-Wallis test: p<0.0001 P-values reflect Mann-Whitney tests compared to *w*. No significant difference was seen between *Myc* and *p53^lh^; Myc*. Related to Figure 2.

**Supplementary Figure 3.**
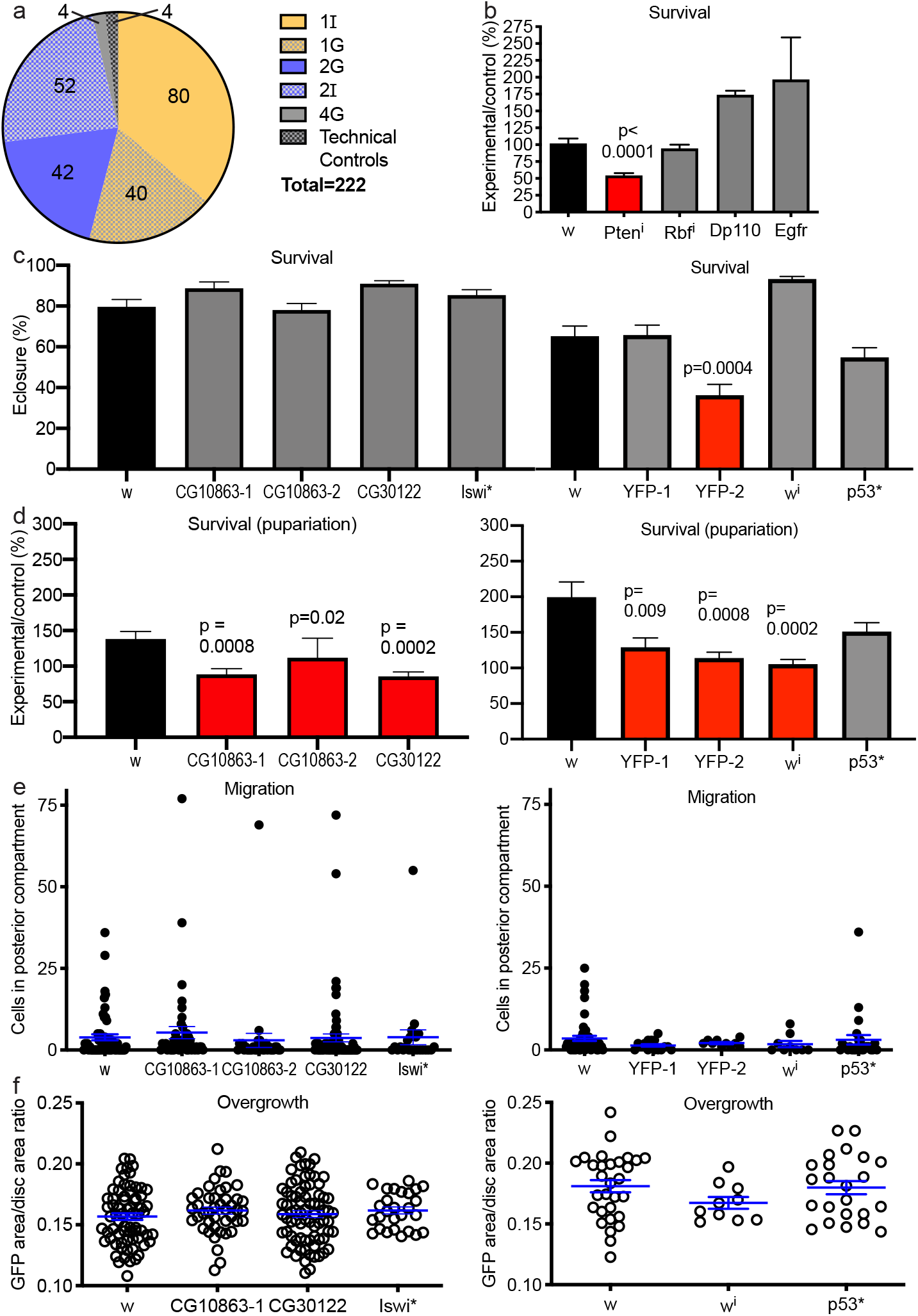
Characterization of positive and negative controls in three *Drosophila* assays. a) Distribution of fly lines assessed in the screen among 6 computationally defined groups. b) Survival of flies to pupariation for positive controls (n=4 for EGFR, n=8 otherwise). c-d) Survival of flies to eclosure (c) and pupariation (d) for low priority genes (left panels: n=8 for CG10863-2 and Iswi*, n=24 for w, n=16 otherwise) and negative controls (right panels: n=8 for all). Pupariation could not be assessed in Iswi* because of a balancer. CG10863-1 and −2 are two lines for the same gene. The pupariation calculation is normalized by counts of internal control pupae and may be more sensitive to noise. e) Quantification of cell migration in Group 4G (left) and control (right) lines. f) Quantification of overgrowth of transgenic tissue in Group 4G (left) and control (right) lines. Overgrowth could not be assessed in CG10863-2 because of a GFP tag, nor the YFP lines. All genotypes shown are in a background of *Myc,p53^sh^*. p-values reflect a student’s t test where data are normally distributed, or a Mann-Whitney test otherwise, compared to *w*. Blue error bars indicate a non-significant difference. Related to Figure 3. See also Supplementary Table 3.

**Supplementary Figure 4.**
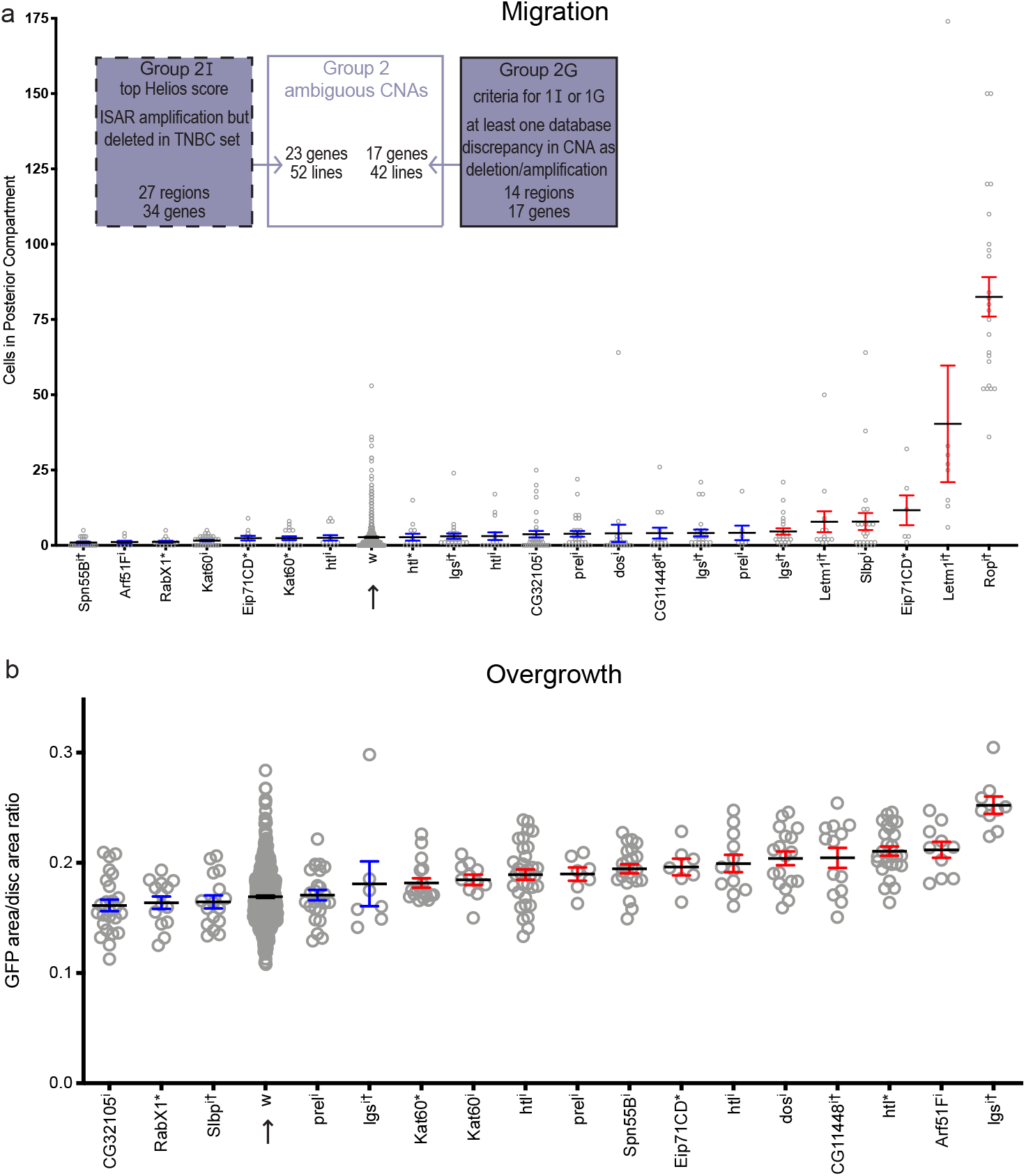
Some driver genes that appear in ambiguous CNAs produce tissue phenotypes in the background of *Myc* and *p53^sh^*. a) Quantification of cell migration for genes from Group 2. Genes marked in red cause significant increase compared to *w* (arrow), measured as p < 0.05 in the original experiment and false discovery rate (fdr) < 0.1 in this aggregate analysis. b) Quantification of transgenic tissue overgrowth for genes from ambiguous deletions. Genes marked in red cause significant increase compared to *w* (arrow), measured asp< 0.05 in the original experiment and fdr <0.1 in this aggregate analysis. Because of variation from one experiment to another, some genes that appear significant in this figure were not significant in their respective experiments. i indicates RNAi against the listed gene; * indicates a heterozygous null allele. † indicates Group 2G; the rest are Group 2I (see Methods). Related to Figure 4.

**Supplementary Table 1**

Results of MutSigCV analysis on TCGA breast cancer somatic mutation dataset.

**Supplementary Table 2**

Computational data on all considered GISTIC 2.0 and ISAR genes.

**Supplementary Table 3**

Fly lines and experimental data for tested genes in Group 4G and negative control genes.

**Supplementary Table 4**

Summary of results for all CNA regions considered in this analysis. Regions in gray are not likely to be significant in TNBC, but may be relevant to other breast cancer subtypes; not all of these were studied to completion.

**Supplementary Table 5**

Data from other databases for all considered GISTIC 2.0 and ISAR genes.

**Supplementary Table 6**

Log-rank test hazard ratios (HR) and p-values for driver genes using the TCGA breast cancer dataset. PFI and OS are preferred by the authors of [39] and are the metrics shown in Figure 5. HR are reported as negative for deletions here, and this is corrected in Figure 5.

**Supplementary Table 7**

Drugs used in the drug screen.

**Supplementary File 1. Fly stocks and results for Group 1 genes**

Results symbols: +: p<0.05. ?: 0.05<p<0.4. -: p>0.4 or rescues phenotype. Each symbol represents one experiment.

**Supplementary File 2. Fly stocks and results for Group 2 genes**

Results symbols: +: p<0.05. ?: 0.05<p<0.4. -: p>0.4 or rescues phenotype. Each symbol represents one experiment.

**Supplementary File 3. Genetic screen results**

In the lethality column, only lines that were significant (p<0.05) in two independent tests are considered positive and reported in each numerator. In the validation column, ‘enhance’ refers to affecting an increase in the phenotype over *Myc,p53^sh^*. Lines were generally only tested for tissue overgrowth (called ‘growth’ in Supplementary Files 1-2) when negative for cell migration; unless otherwise indicated, a test for tissue overgrowth implies a negative cell migration test. ‘Passenger’ refers to genes lacking functional evidence for driver status in this study. ND = not done.

**Supplementary File 4. Migration assay results**

Stock numbers and statistical results corresponding to Figures 4a and S4a.

**Supplementary File 5. Overgrowth assay results**

Stock numbers and statistical results corresponding to Figures 4b and S4b.

